# Construction of synthetic microbiota for reproducible flavor metabolism in Chinese light aroma type liquor produced by solid-state fermentation

**DOI:** 10.1101/510610

**Authors:** Shilei Wang, Qun Wu, Yao Nie, Yan Xu

**Affiliations:** State Key Laboratory of Food Science and Technology, Synergetic Innovation Center of Food Safety and Nutrition, School of Biotechnology, Jiangnan University, Wuxi, Jiangsu, 214122, China; Suqian Industrial Technology Research Institute of Jiangnan University, Suqian, Jiangsu, 223800, China

**Keywords:** Chinese liquor, Co-occurring network, Core microbiota, Environmental factors, Flavor compounds, Food fermentation

## Abstract

Natural microbiota plays an essential role in flavor compounds producing for traditional food fermentation. Whereas, the fluctuation of natural microbiota results in the inconstancy of food quality. Thus, it is critical to reveal the core microbiota for flavor compounds producing and construct a synthetic core microbiota for constant food fermentation. Here, we revealed the core microbiota based on their flavor-producing and co-occurrence performance, using Chinese light aroma type liquor as a model system. Five genera were identified to be the core microbiota, including *Lactobacillus, Saccharomyces, Pichia, Geotrichum*, and *Candida*. The synthetic core microbiota of these five genera presented a reproducible dynamic profile with that in the natural microbiota. Monte Carlo test showed that the interpretation of five environmental factors (lactic acid, ethanol and acetic acid contents, moisture and pH) on the synthetic microbiota distribution were highly significant (*P* < 0.01), which was similar with that in the natural fermentation system. In addition, 77.27% of the flavor compounds produced by the synthetic core microbiota showed a similar dynamic profile (*ρ* > 0) with that in the natural liquor fermentation process, and the flavor profile presented a similar composition. It indicated that the synthetic core microbiota is efficient for reproducible flavor metabolism. This work established a method for identifying core microbiota and constructing a synthetic microbiota for reproducible flavor compounds. It is of great significance for the tractable and constant production of various fermented foods.

**IMPORTANCE:** The transformation from natural fermentation to synthetic fermentation is essential to construct a constant food fermentation process, which is the premise for stably making high-quality food. According to the functions of flavor-producing and co-occurring in dominant microbes, we provided a system-level approach to identify the core microbiota in Chinese light aroma type liquor fermentation. In addition, we successfully constructed a synthetic core microbiota to simulate the microbial community succession and flavor compounds production in the *in vitro* system. The constructed synthetic core microbiota could not only facilitate a mechanistic understanding of the structure and function of the microbiota, but also be beneficial for constructing a tractable and reproducible food fermentation process.

## INTRODUCTION

Traditional fermented foods are usually produced by natural fermentation containing multi-species community (1-4). At present, the transformation from natural fermentation to tractable fermentation with the synthetic core microbiota is essential for consistent quality of fermented foods because only limited genera of microbes in natural microbiota can drive the fermentation process. They not only generate flavor compounds but also maintain microbes’ interaction, which serve to achieve the successful food fermentation (5, 6). Thus, revealing the composition of these microbes, that is the core microbiota, is essential for constructing a synthetic microbiota in food fermentation (7).

A series of studies were carried out to identify the core microbiota during food fermentation (7-10). Dominant genera were considered to be an essential component in the food fermentation (11, 12). For example, a total of 17 genera were identified to be dominant microbes due to their relative abundance in cheese (11). However, dominant genera may not have the ability to produce flavor compounds in food fermentation (13, 14). Researchers suggested that the identification of core microbiota should also consider the microbial flavor compounds productivity (7, 15). For example, seven genera were determined as functional core microbiota for production of flavor compounds in Chinese vinegar fermentation (7).

Recently, we found that only the dominant microbes or flavor-producing microbes did not show efficient flavor compounds productivity when they were in a mixed culture (13). Whereas, some other microbes were not flavor compound producers, but they showed the activity to coordinate with those flavor-producing microbes, hence leading to an improvement of flavor compounds (13). For example, *Pichia membranaefaciens* and *Bacillus amyloliquefaciens* were not efficient flavor compound producers, but they alleviated the competition among flavor compound producers (*Saccharomyces cerevisiae, Issatchenkia orientalis* and *Bacillus licheniformis*), and finally altered the producers’ growth and flavor compound productions (13). Moreover, the interaction between microbes plays a vital role in some flavor metabolisms, such as 3-(methylthio)-1-propanol and dimethyl disulfide (16). As a consequence, we suggest that besides the flavor compounds productivity, the microbial interaction should also be considered to identify the core microbiota. Moreover, microbial interaction is a critical factor for maintaining the co-occurring in microbial communities, and co-occurring network analysis is an effective tool for studying the microbial interaction (17, 18).

Thus, to overcome the problem of inaccurate definition of core microbiota in fermented foods, we provided a comprehensive method to identify the core microbiota in natural food fermentation, with the combination of flavor-production and co-occurring network analysis. We also took a prudent way to examine the activity of the core microbiota, including its interaction with the environmental factors, and the flavor compound producing. Due to the Chinese light aroma type liquor is a favorite alcoholic beverage and generated by a natural fermentation process (19). In this work, using Chinese light aroma type liquor fermentation as a model system, we provided a strategy to identify the core microbiota and constructed a synthetic microbiota using the core microbiota. Because, Chinese light aroma type liquor, a typical and popular fermented food, is made from spontaneous fermentation involving multiple microbes and complex interactions between microbes (12, 20), and this type of fermentation can produce unique food flavor and taste characteristics (1). And it is also one of the three typical type liquors in China (Sauce aroma, Strong aroma, and light aroma type liquor). In addition, it has a smaller brewing container, shorter fermentation time and easy to observe. So, taking the Chinese liquor as a model system and establishing a method to define the core microbiota are beneficial for constructing a synthetic microbiota to reveal the mechanism of fermented foods.

## RESULTS

### Microbial diversity during the fermentation process

Across all samples, altogether 453,217 and 677,563 high-quality sequences were identified for bacteria and fungi after quality control. Meanwhile, a total of 722 and 1,504 operational taxonomic units (OTUs) were obtained for bacteria and fungi with 97% similarity. A total of 49 bacterial genera and 34 fungal genera were identified in the fermentation process (Dataset S1). All the Good’s coverage of samples were over 99.80% (Table S1) that indicated sequences represented the majority of microbiota in the fermentation process (21). The average bacterial α-diversity (Chao1 richness and Shannon diversity) declined along with fermentation time on the whole, but there was a fluctuation on day 15 (Table S1). On the contrary, the average fungi α-diversity (Chao1 richness and Shannon diversity) increased along with fermentation time on the whole, but there was a fluctuation on day 5 (Table S1).

As for bacteria (Fig. 1A), at the early stage of fermentation (0 day), Pseudomonas and Bacillus were the predominant genera (average abundance ≥ 10%) (22), whereas *Lactobacillus*, *Pediococcus*, *Leuconostoc*, *Weissella*, *Stenotrophomonas*, *Staphylococcus*, *Streptomyces*, *Kroppenstedtia*, *Herbaspirillum*, *Achromobacter*, *Flavobacterium* and *Brevibacterium* were subdominant genera (1% ≤ average abundance ≤ 10%). During middle stage of fermentation (day 5-15), *Lactobacillus* and *Pediococcus* became the predominant genera, and *Leuconostoc* was the subdominant genus at day 15. At the late stage of fermentation (day 20-28), only *Lactobacillus* was the predominant genus, *Pediococcus* became the subdominant genus. As for fungi (Fig. 1B), *Pichia* was the predominant genus in the whole fermentation process. *Geotrichum* (day 5-10) and *Saccharomyces* (day 10) were the predominant genera, and *Saccharomycopsis*, *Rhizopus*, *Clavispora*, *Candida*, *Aspergillus*, *Thermomyces, Thermoascus, Trichosporon*, and *Lichtheimia* were the subdominant genera at different stages of fermentation.

**Fig. 1.**
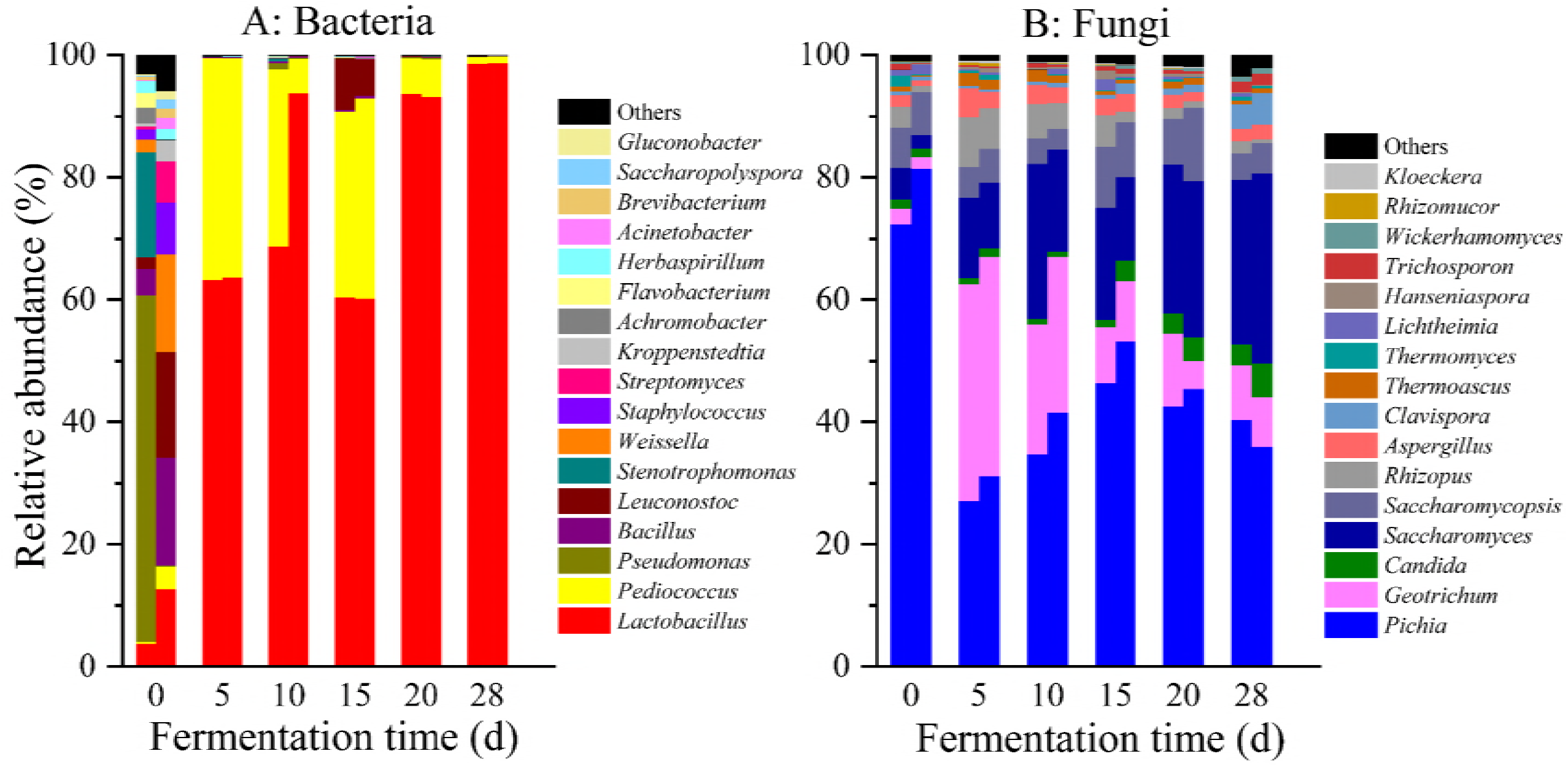
Distribution of the relative abundance of genera during the fermentation in the *in situ* System. Only those genera that had an average abundance greater than 1% were indicated. Genera less than 1% abundance were combined and shown in others.

Through statistical analysis of all communities sampled, only 17 bacterial and 16 fungal genera were found at greater than 1% average abundance, which were defined as dominant microbiota (9). A ubiquitously distributed microbiota is usually defined as being present in most samples (9, 23). Therefore, we defined microbes, which exist in more than 50% samples within a total of 14 samples, as ubiquitously distributed microbiota (9). Two genera of bacteria (*Lactobacillus* and *Pediococcus*) and eight genera of fungi (*Pichia*, *Geotrichum*, *Saccharomyces, Saccharomycopsis, Rhizopus, Aspergillus, Candida* and *Thermoascus*) were identified to be ubiquitously distributed dominant microbiota (Table S2).

### Identification of the core microbiota

Flavor compounds are very important indicators of the liquor quality (24, 25). A total of 41 kinds of flavor compounds were identified during the fermentation (Fig. 2), including four alcohols, two carbonyl compounds, five acids, 20 esters, nine aromatic compounds and one heterocyclic compound.

**Fig. 2.**
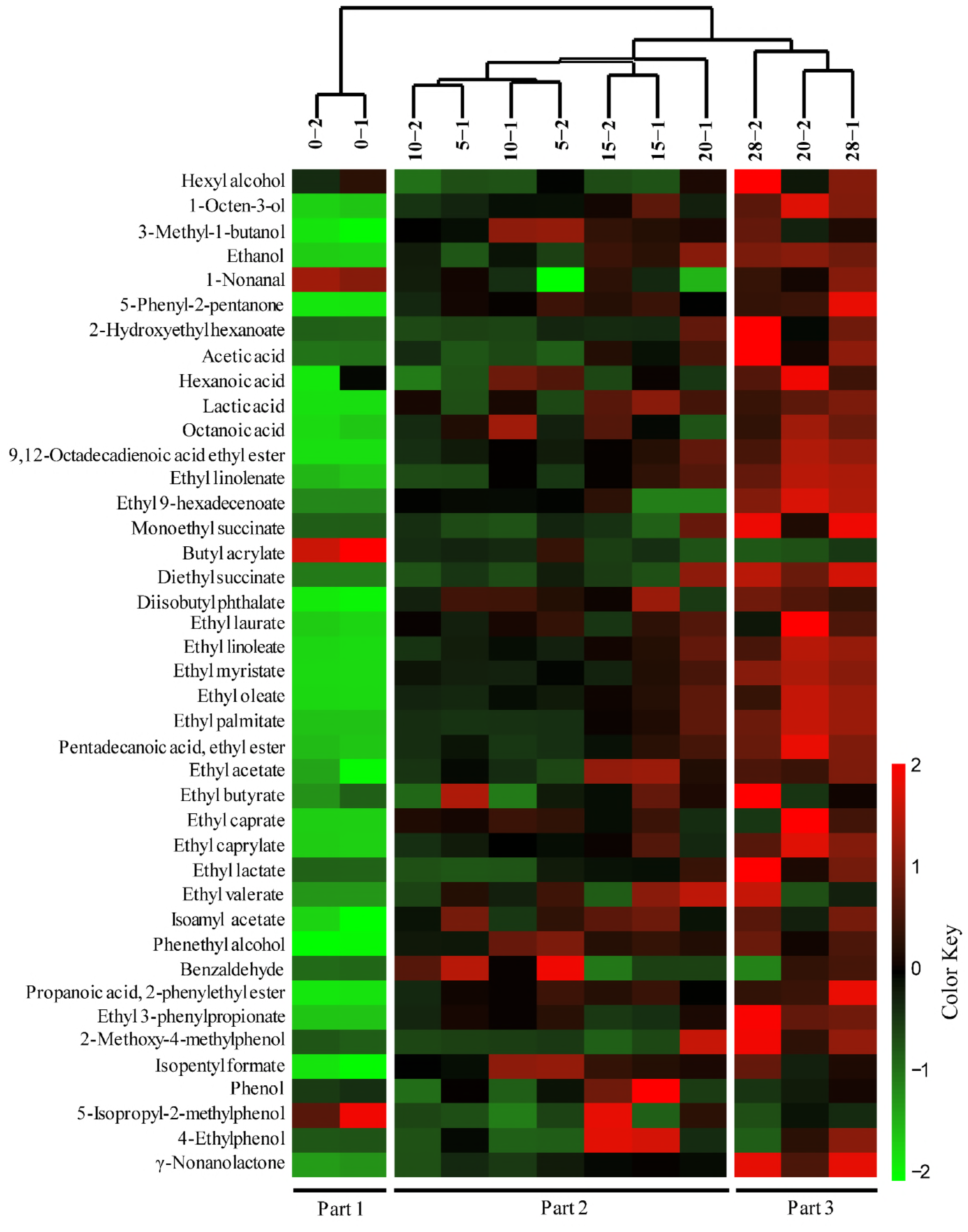
Heatmap of flavor metabolites and hierarchical clustering in the *in situ* fermentation process. Flavor compounds were transformed by z-score. Clustering analysis was performed using the Pearson correlation coefficient and Euclidean distance based on the flavor contents during the fermentation process.

Concentrations of flavor compounds were transformed and converted into a heat map, and hierarchical cluster analysis was achieved. As shown in Fig. 2, the hierarchical clustering results showed that the fermentation process consisted of three-part based on the dynamic profile of flavor compounds: part 1 (day 0), part 2 (days 5-20) and part 3 (days 20-28).

Most flavor compounds are related to microbes in food fermentation. Network correlation analysis is a powerful tool to investigate the potential interactions between microbes and flavor compounds (26). Thus, we calculated the Spearman correlation coefficient between 33 dominant genera and 41 flavor compounds, and chose the coefficient (*ρ*) > 0.5 and significance (*P*) < 0.05 (27, 28) as strongly correlated nodes of the network (Fig. 3 A). Eight bacterial and seven fungal genera were significantly correlated (*P* < 0.05, *ρ* > 0.5) with 34 kinds of flavor compounds, indicating these 15 genera are the flavor-producing microbiota (Table S3). Among them, *Lactobacillus, Saccharomyces, Clavispora* and *Candida* were significantly correlated (*P* < 0.05, *ρ* > 0.5) with 26, 26, 16 and 14 kinds of flavor compounds, respectively (Fig. 3 A).

**Fig. 3.**
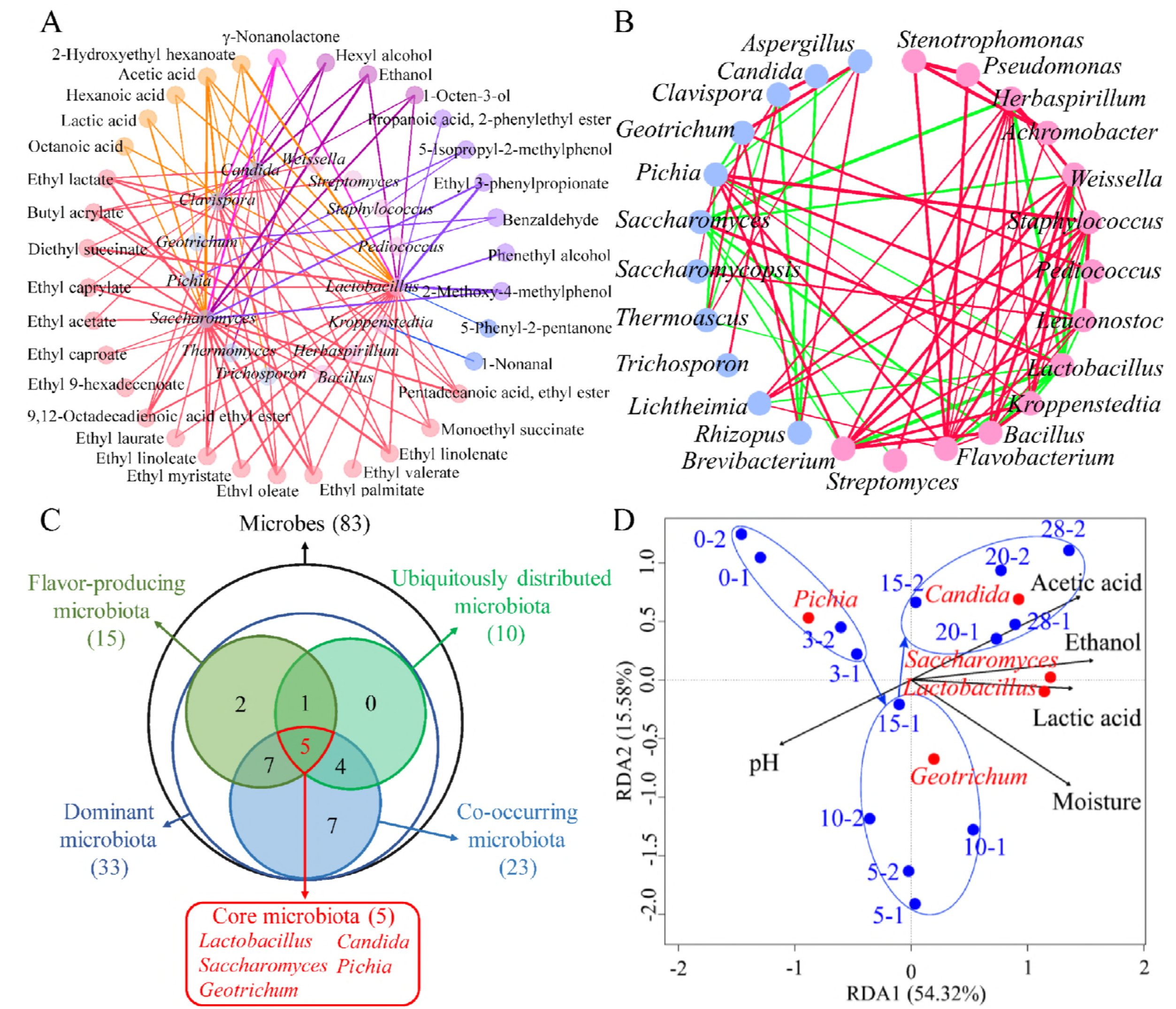
Identification of the core microbiota in the *in situ* system. (A) Correlation network between microbial genera and flavor compounds during the fermentation process in the *in situ* system. Inner circle nodes represent microbes (light red modules represent bacteria genera, and light blue modules represent the fungi genera), and outer circle nodes represent flavor compounds (different colors represent different flavor types). The thickness of lines are proportional to the value of Spearman’s correlation (*ρ* > 0.5, *P* < 0.05). The color of lines are same with the flavor nodes. (B) Correlation network of co-occurring genera in dominant microbiota. Statistically significant (*P* < 0.05) and Spearman correlation coefficient (|*ρ*| > 0.5) indicate the correlations. Light red modules represent bacteria genera, and light blue modules represent fungi genera. Green and red edges indicate negative and positive interaction between genera. The thickness of lines represents the strength of interaction. (C) The Venn diagram of the core microbiota. Different circles represent different genera categories. (D) RDA analysis of fermentation process. Blue dots represent the time of fermentation. Red dots represent the core microbiota. Black arrows represent the different of environmental factors. Percentages on the axis represent the eigenvalues of principal components.

Co-occurrence network analysis allows identifying the co-occurring microbiota (17). We calculated the Spearman correlation coefficient of 33 dominant genera. The Spearman’s correlation coefficient (|*ρ*|) > 0.5 and significance (*P*) < 0.05 was considered to be a valid co-occurrence event (17, 18, 26, 29, 30). Through the co-occurrence network analysis, a total of 25 nodes and 149 edges were obtained (|*ρ*| > 0.5, *P* < 0.05), and the average network clustering coefficient was 0.696, which suggested that the network had modular structures. In Fig. 3 B, different genera are divided into different modular structures. A total of 23 genera presented highly connection (≥ 4 edges per node) (26), and were defined as the co-occurring microbiota, including *Flavobacterium, Lactobacillus*, *Brevibacterium*, *Herbaspirillum*, *Pichia, Staphylococcus, Bacillus, Weissella, Kroppenstedtia, Leuconostoc, Saccharomyces, Aspergillus*, *Clavispora*, *Geotrichum*, *Lichtheimia*, *Thermoascus*, *Rhizopus*, *Achromobacter*, *Pseudomonas*, *Stenotrophomonas*, *Candida*, *Saccharomycopsis* and *Streptomyces* (Table S4). In the co-occurrence network, *Lactobacillus* and *Saccharomyces* were mainly negatively correlated (*ρ* < - 0.5) with other microbes (excepting *Clavispora*), but they showed a positive correlation with each other.

In summary, we obtained ubiquitously distributed dominant microbiota (10 genera), flavor-producing microbiota (15 genera) and co-occurring microbiota (23 genera). Five genera existed in all these three different microbiotas, including *Lactobacillus, Saccharomyces*, *Geotrichum*, *Candida* and *Pichia* (Fig. 3 C). Due to their high relative abundance and frequency, the contributions to flavor productions and the stable microbial network, they were defined as core microbiota in liquor fermentation.

The impact of five environmental factors on the core microbiota were analyzed, including lactic acid content, ethanol content, acetic acid content, moisture and pH (Table S5). Variation partitioning analysis (31) was used to calculate the contributions of these environmental factors. The results showed that these five environmental factors accounted for 87.18% of core microbiota’s variation in the *in situ* systems (Table S6). Partial redundancy analysis (RDA) was used to identify the effect of these factors on the core microbiota (Fig. 3 D). Acetic acid content, ethanol content, and lactic acid content were positively correlated with *Lactobacillus, Saccharomyces* and *Candida* at the end of fermentation. Monte Carlo replacement test (Table S7) verified the result that these factors were significantly correlated with the core microbiota (*P* < 0.05). It indicated these five environmental factors had a significant influence on the core microbiota.

### Reproducible dynamic profile of microbiota in synthetic core microbiota

In this study, we provided a system-level approach to identify the core microbiota in Chinese light aroma type liquor fermentation, and obtained five different core genera during the whole fermentation stage, including *Lactobacillus*, *Pichia, Geotrichum*, *Candida* and *Saccharomyces*. Due to the diversity of genera, it was considered to be feasible that isolated species represented certain taxa. For example, cheese rind isolates that represented the most abundant taxa were applied to construct in vitro communities of cheese rind (9). Using 16S and ITS amplification sequence data, when the sequence identity was greater than 99% compared to the type and reference strains, the assignment to the species level was performed (32). Thus, we identified one species with the highest relative abundance in each corresponding genera (Fig. S1 and Table S8) and used as the starter species of the synthetic microbiota, including *Lactobacillus acetotolerans*, *Pichia kudriavzevii*, *Geotrichum candidum, Candida vini* and *Saccharomyces cerevisiae. Lactobacillus acetotolerans* is a functional microorganism in the fermentation of kinds of liquors (Strong aroma, light aroma type liquor and Japanese sake) (32-34). For example, *Lactobacillus acetotolerans* appeared to play a key role during the Chinese strong aroma type liquor fermentation (32), and it had positive relationships with most chemical components that contribute to the quality and flavor of liquor (35). *Pichia kudriavzevii* contribute to the functionality (acids and esters) of foods during fermentation, and it can improve the sensory and some functional properties of the cereal-based substrate during fermentation (36). *Geotrichum candidum*, can produce lipases, which would be important for the productions of fruity aroma compounds (37). *Candida vini* had been shown to contribute to fatty acids (38). Saccharomyces cerevisiae is an important strain of ethanol fermentation in Chinese liquor fermentation (39). Therefore, we chose the five species for the synthetic experiment.

We inoculated approximately equal numbers of each species in the five core genera together into fermented grains in the *in vitro* system (Fig. 4 A). *Lactobacillus* became the predominant genus in the *in vitro* system as fermentation proceeded (Fig. 4 A and Fig. S2 B), which was similar with that of the *in situ* system (Fig. S2 A, Fig. S3). *Saccharomyces* and *Pichia* were the dominant genera in the early (1-5 d) and end (28 d) of the fermentation process, which was similar with that of the *in situ* system (Fig. 4 A, Fig. S2 C and Fig. S2 D). *Candida* was the dominant genera in the middle fermentation process (10-25 d). It revealed that the successive direction of the *in vitro* systems (Fig. 4 B) in the principal component is consistent with that of the *in situ* system over a 28 d fermentation period (Fig. 4 C), which demonstrated a highly reproducible microbial succession pattern of the *in vitro* liquor fermentation.

**Fig. 4.**
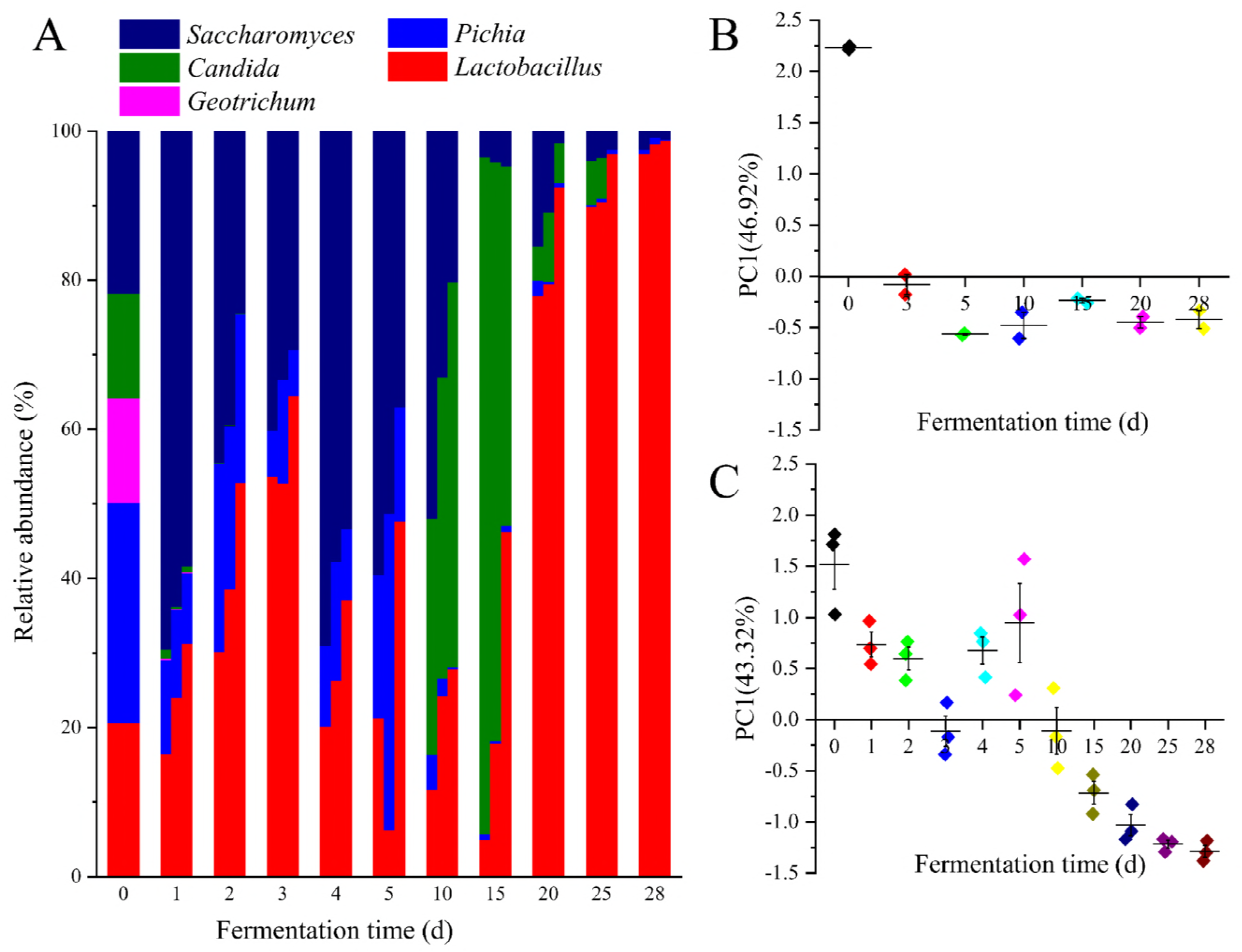
Reproducible dynamic profile of microbiota in synthetic core microbiota. (A) Distribution of the abundance of genera during the fermentation in the *in vitro* system. (B) *In situ*, the change of principal component in time gradient. (C) *In vitro*, the change of principal component in time gradient.

The impact of the environmental factors on the synthetic microbiota were also analyzed (Table S9). Explanations of variation partitioning analysis of the five environmental factors (lactic acid content, ethanol content, acetic acid content, moisture and pH) reached 53.65% in the *in vitro* system (Table S6). This percentage showed that these five factors drove the variation of the synthetic core microbiota. RDA analysis showed that pH was negatively correlated with the other environmental factors that was the same with that in the *in situ* system (Fig. 5 A). Lactic acid content, acetic acid content, ethanol content and moisture were positively correlated with each other that are consistent with that of the *in situ* system. Monte Carlo test also showed that the interpretation of these five environmental factors on the synthetic microbiota distribution were highly significant (*P* < 0.01) (Table S7). Through the change of environmental factors’ correlation analysis of the two systems on the temporal dynamics (Fig. 5 B), we found that five environmental factors had a positive correlation (*ρ* > 0) with the core microbiota, especially, moisture, acetic acid content, lactic acid content and pH had a strong correlation (*ρ* > 0.6) between in the *in situ* and *in vitro* systems. These results indicated that the effect of environmental factors on the core microbiota was also similar in the *in vitro* and *in situ* systems.

**Fig. 5.**
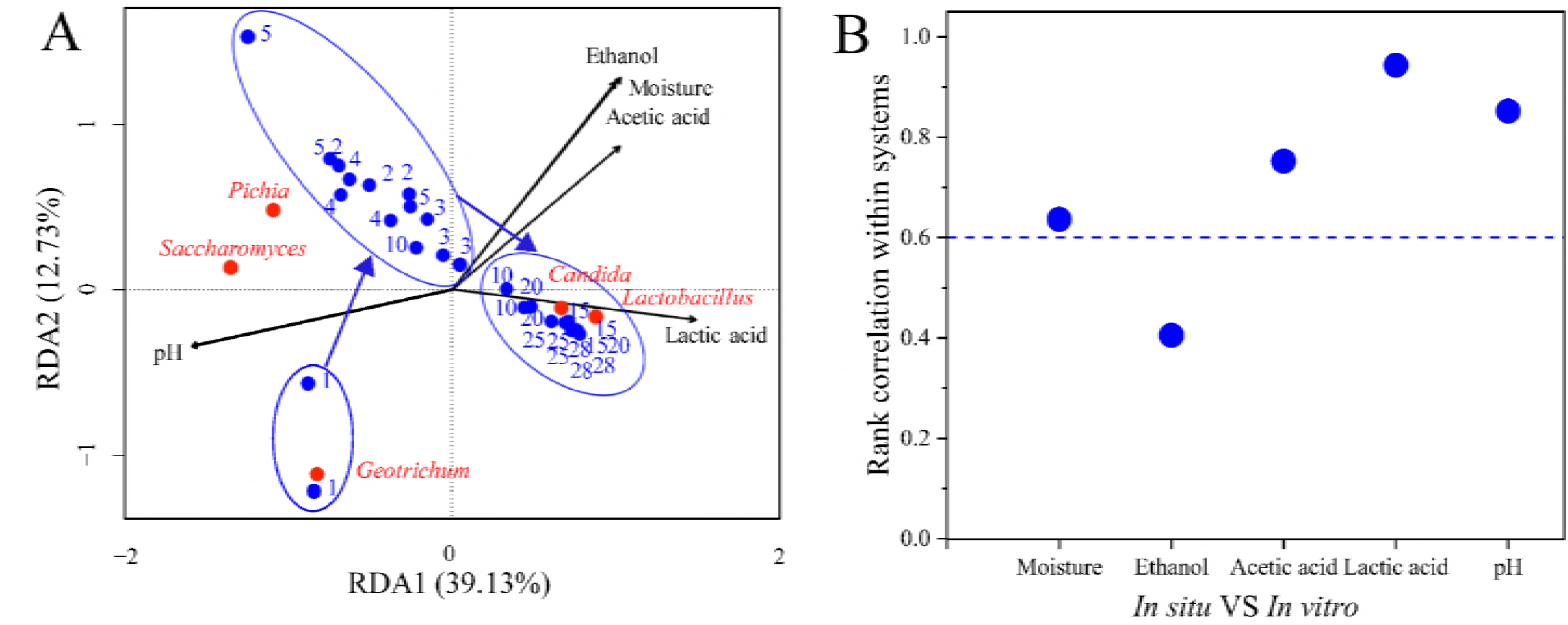
RDA analysis of fermentation process in the *in vitro* system and the relationship of environmental factors within the *in situ* system. (A) RDA analysis of fermentation process in the *in vitro* system (same with Fig. 3D). (B) The similarity of in the *in situ* and *in vitro* system. The vertical coordinate in the figure represents the Spearman correlation coefficient between the corresponding of environmental factors in the two systems.

### Reproducible flavor metabolism in synthetic core microbiota

The flavor compound producing in the synthetic microbiota was determined, and 22 flavor compounds were identified in the *in situ* system (Fig. 6 A). The *in vitro* generation of flavor compounds can be divided into three parts (Fig. S4): part 1 (day 0-3), part 2 (day 4-10), part 3 (day 15-28). The temporal dynamics was similar to that in the *in situ* system.

**Fig. 6.**
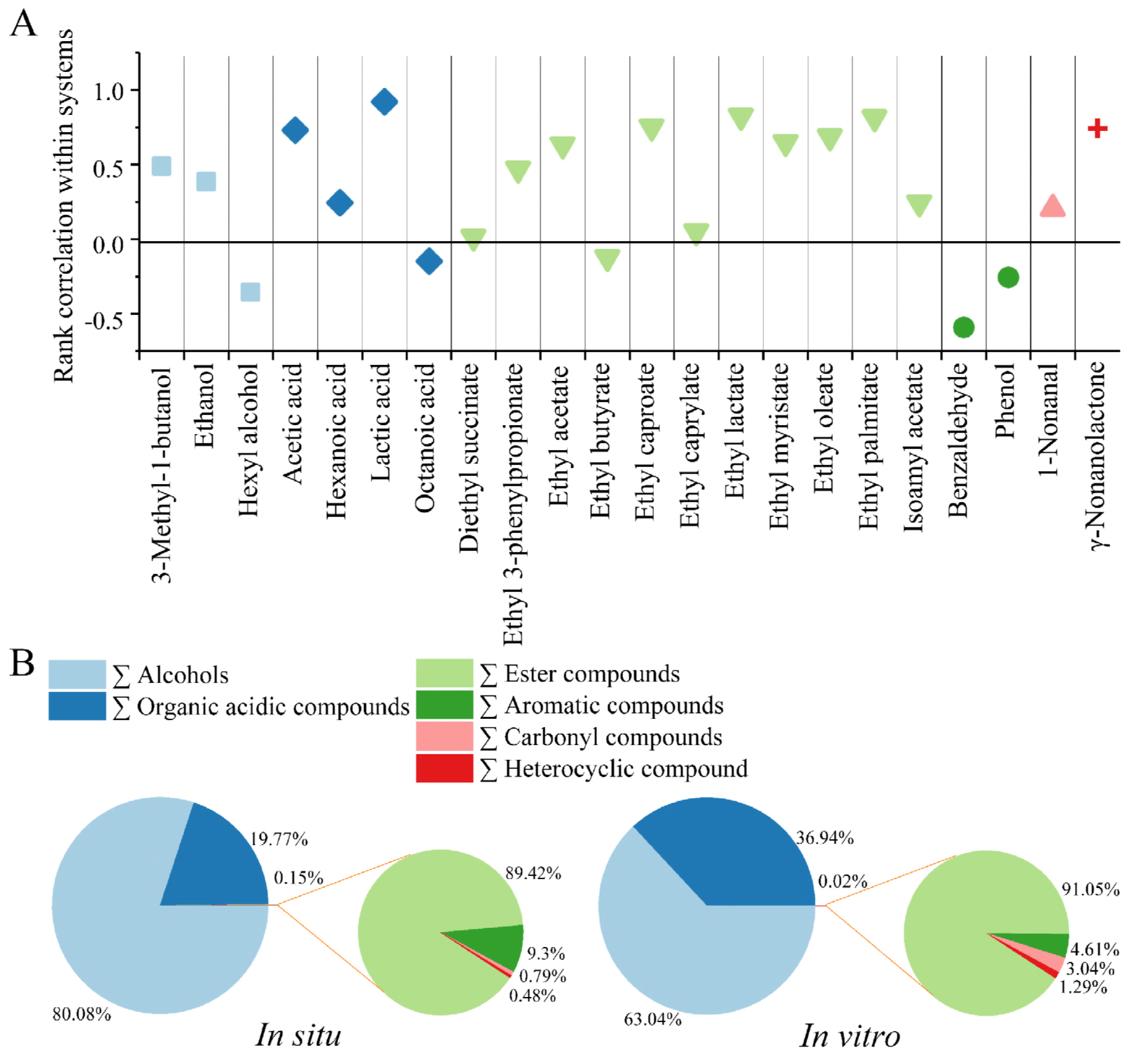
Reproducible flavor metabolism in synthetic core microbiota. (A) The similarity of two system in 22 kinds of alcohols, acids and esters in two systems. The vertical coordinate in the figure represents the Spearman correlation coefficient (*ρ*) of the flavor generation along the time axis in the two systems. (B) The ratio of six kinds of flavor compounds in the *in situ* and *in vitro* systems.

The Spearman correlation coefficient (*ρ*) of the 22 flavor compounds generation in the two systems was calculated in the fermentation. The result showed that 17 kinds of flavor compounds (ratio = 77.27%) had a positive correlation (*ρ* > 0) with the generation on the temporal dynamics in the two systems (Fig. 6 A). The ratio of different flavor classifications had similar proportions in both systems (Fig. 6 B), in which the ratio of alcohols and acids accounted for more than 99.85% in the total flavor compounds. These indicated that the flavor metabolism could be reproduced in the *in vitro* system with the synthetic core microbiota.

## DISCUSSION

Core microbiota inhabiting in food fermentation is of great importance to the quality and characteristics of foods. Many molecular and ecological approaches have been used to characterize the core microbiota (22, 40-42). In this work, we chose microbial communities in Chinese light aroma type liquor fermentation as a model system and provided a system-level method to identify the core microbiota in natural food fermentation. Taking a prudent way examined the characteristics of all the dynamic succession of the microbiota, the effect of the environmental factors, and the profile of flavor compounds production. Among these compounds, we did not detect the detrimental flavors in Chinese light aroma type liquor fermentation. Most of these flavors have pleasant aromatic smells, such as ethyl acetate (pineapple), ethyl lactate (fruity), 1-Octen-3-ol (mushroom), octanoic acid (cheesy), ethyl 3-phenylpropanoate (floral), γ-nonanolactone (coconut), etc. (43, 44). Although some of these flavors also contain some unpleasant flavors, but these flavors form a special style of products at low concentrations, such as acetic acid (acidic, vinegar), hexanoic acid (sweaty), ethyl oleate (fatty), 3-methyl-1-butanol (malty), etc. (43, 44). We constructed a reproducible synthetic core microbiota, with that of the natural microbiota for liquor fermentation. It would help us to establish a tractable food fermentation system.

In the *in vitro* system, alcohols (without ethanol) and acid contents were a bit higher than those in the *in situ* system (Dataset S2, Dataset S3). Whereas, ester contents were lower than that in the *in situ* system (*P* < 0.001). That may be due to the low concentration of esterification strains in the *in vitro* system. We also observed slight differences in the microbiota between the *in situ* and *in vitro* systems. For example, a succession of *Saccharomyces* appeared to proceed much more quickly (Fig. S2 C, S2 D), and *Candida* showed a higher relative abundance in the later fermentation in the *in vitro* system (Fig. 4, Fig. S2 D). The difference might result from a higher initial ratio of these genera in the *in vitro* system. Therefore, the initial compositions of the core microbiota should be optimized in the further synthetic core microbiota’s fermentation. Different species and different strains of microorganisms under the same genus may have different metabolic functions. Therefore, more functional strains should be isolated. But the same strain in single fermentation and mixed fermentation may show completely different metabolic patterns (16). Therefore, the target functional strains should be synthetically optimized by extensive statistical analysis.

Besides the liquor fermentation system, the methods for identifying the core microbiota and constructing a synthetic microbiota for food fermentation can also be used in a variety of food fermentation processes. Various food fermentations share the members of the core microbiota because these members present similar functions in different food fermentations. For example, *Lactobacillus* was confirmed to be the core microbe in fermentations of vinegar, liqueur, cheese, pickle, and so on (45-47). It contributed amino acids (glutamic acid, alanine, valine, etc.), organic acids (acetic acid, lactic acid, etc.), and other flavor compounds (7, 48-50). It would also interact with other microbes, such as *Bacillus, Aspergillus* and *Luteococcus*, hence regulate their flavor compounds producing (47, 50-52). *Pichia* was widely used in food fermentation, such as wine and beer fermentation (53-55). It was considered to be an essential producer of esters (56). *Pichia* can also maintain the co-occurring of the community (13), which was similar to that in Fig 3 B. *Geotrichum* can produce lipases, which would be important for the productions of fruity aroma compounds, such as ethyl esters of acetic acid, propionic acid, butyric acid and isobutyric acid (57, 58). *Saccharomyces*, as an ethanol producer, was widely used in liquor and other alcoholic beverages’ production (59). It drove the development direction of the microbiota, together with *Lactobacillus* (acid producer) (60, 61). *Candida* was widely used in food fermentations, due to its production of various lipases (antarctica lipase A, rugosa lipases, glucose ester synthesis lipase, *etc*.) (62-64). When *Candida* and *Saccharomyces* were co-cultured in wine fermentation, they produced higher amounts of esters and glycerol, compared with that of single *Saccharomyces* (65). These studies indicated that most of the microbes in the core microbiota had similar functions in different food fermentations.

The transformation from natural fermentation to synthetic fermentation is essential to construct a tractable food fermentation process, which is the premise for stably making high-quality foods. We provided a system-level approach to identify the core microbiota in food fermentation and constructed a synthetic microbiota for reproducible flavor metabolism. It would provide a chance for us to define the mechanisms underlying the microbial interaction and contribution to flavor compounds in the food microbiota. It is also important to manipulate the synthetic microbiota and then control the quality of fermented foods.

## MATERIALS AND METHODS

### Sample collection

Samples were collected from a local liquor distillery (Shanxi Xinghuacun Fenjiu Distillery Co. Ltd. Shanxi, China,). For liquor fermentation, the steamed grains were mixed with starter at a ratio of 9:1 (w/w) and put into earthenware jars. Then, the jars were sealed for 28 days’ fermentation. For the survey of microbial diversity, a total of 12 samples (100 g each sample) were collected from 2 jars in the center of the layer (0.5 m deep) at different fermentation times (day 0, 5, 10, 15, 20, and 28) in April 2016. All samples were stored at - 20°C for further DNA extraction and physicochemical parameters determination.

### DNA extraction, qualification and sequencing analysis

Each sample (5.00 g) was used to extract genomic DNA using the E.Z.N.A. ^®^ soil DNA Kit (Omega Bio-tek, Norcross, GA) according to manufacturer’s instruction. The V3-V4 region of the 16S rRNA bacterial gene was amplified with the universal primers 338F (5’-ACTCCTACGGGAGGCAGCAG-3) and 806R (5’-GACTACHVGGGTWTCTAAT-3’) (66). For fungi, the ITS2 region was amplified with the primers of ITS2 (5’-GCTGCGTTCTTCATCGATGC -3’) and ITS3 (5’-GCATCGATGAAGAACGCAGC -3’) (67). These primers added a set of 8-nucleotide barcodes sequence unique to each sample. The PCR reactions were performed in 25 μL volume, containing 2.5 μL of 10 × Pyrobest Buffer, 2 μL of 2.5 mM dNTPs, 1 μL of each primer (10 μM), 0.4 U of Pyrobest DNA Polymerase (TaKaRa, Takara Holdings Inc., Nojihigashi, Kusatsu, Shiga, Japan), 15 ng of template DNA, and double-distilled water (ddH_2_O) up to 25 μL. Amplification was performed with the previously described method (42, 68). Then applicants were pooled into equimolar quantities and subjected to high-throughput sequencing using Miseq Benchtop Sequencer for 2 × 300 bp pair-end sequencing (Illumina, San Diego, CA). Databases of EzBioCloud and Central Bureau of Fungal Cultures (CBS-KNAW) were used for sequence alignment of bacteria and fungus. The fungi and bacterial raw sequences data were deposited in the DNA Data Bank of Japan (DDBJ) database under the accession number of DRA005471 and DRA005916.

### Population determination by real-time quantitative PCR (qPCR)

The population of yeast and LAB in liquor fermentation were determined by qPCR. Genomic DNA of samples were used as the templates. For yeast, the sequences were amplified using YEASTF (5’ GAGTCGAGTTGTTTGGGAATGC 3’) and YEASTR (5’ TCTCTTTTCCAAAGTTCTTTTCATCTTT 3’) (69) as primers. For LAB, the sequences were amplified using Lac1 (5; AGCAGTAGGGAATCTTCCA 3’) and Lac2 (5’ ATTYCACCGCTACACATG 3’) (70) as primers. qPCR was performed by the StepOnePlus instrument (Applied Biosystems, CA, USA) (16).

### Sequence processing

All the raw Miseq-generated sequences were processed via QIIME (V. 1.8) (71). Briefly, high-quality sequences was carried out by removing sequences with ambiguous bases > 2, homopolymers > 10, primer mismatches, average quality scores < 20 and lengths (excluding the primer or barcode region) < 50 bp. Chimeras were removed using USEARCH (v. 10) (72). The trimmed sequences were clustered into operational taxonomic units (OTUs) with 97% sequence (73), and then calculated Shannon index and Chao1 estimator using UCLUST (V. 1.2.22) (74, 75).

### Analysis of environmental factors and flavor compounds

Moisture was measured by determining its weight loss after drying 10 g of each sample at 105°C for 3 h (sufficient to ensure constant weight) (76). The pH was measured at a 1:2.5 (w/v) ratio in distillation-distillation H_2_O (ddH_2_O) with the Laboratory pH meter-FE20 (Mettler Toledo, Shanghai, China) (76). Five-gram samples were added to 10 mL ddH_2_O and put in an ultrasonic cleaner (AS30600B, Autoscience, Tianjin, China) for 30 min, and then centrifuged at 8000 × *g* for 10 min. After filtered using a 0.2 μm filter, the filtrate was used to analyze the concentrations of flavor compounds and acids. The flavor compounds content was detected using gas chromatography-mass spectrometry (Agilent 6890N GC system and Agilent 5975 mass selective detector, Agilent, Santa Clara, CA) (42). The ethanol content was determined by high-performance liquid chromatography (HPLC, Agilent 1200, Agilent, Santa Clara, CA) using a column aminex HPX-87H (Bio-Rad, Hercules, CA) (77). The contents of lactic acid and acetic acid were measured using reversed-phase ultra-performance liquid chromatography (UPLC, Waters H-class system, Waters, Milford, MA) with chromatographic column waters Atlantis T3 (4.6 mm×150 mm, 3 μm) (Waters, Milford, MA) and guard column Phenomenex RP-C_18_ Security Guard (4.0 mm×3.0 mm) (Phenomenex Inc. Torrance, CA). The UV detection wavelength was 210 nm. The column temperature was 30°C. The injection volume was 10 μL. The mobile phase was 10 mmol/L NaH_2_PO_4_ (pH 2.7), and the flow velocity was 0.8 mL/min.

### Strains

Predominant microbes were all isolated from the liquor fermentation process, *Lactobacillus acetotolerans*, *Pichia kudriavzevii* and *Candida vini* were deposited in China General Microbiological Culture Collection Center with the accession number of CGMCC No. 14086 and 12418 and 2.2018. *Saccharomyces cerevisiae* was deposited in China Center for Type Culture Collection with the accession number of CCTCC M2014463. *Geotrichum candidum* is a laboratory strain with the number of XY7.

### Liquid fermentation

The sorghum extract was used as seed fermentation broth (40). The extract was diluted with distilled water to give a sugar concentration of about 90 g/L and then autoclaved at 115°C for 15 min. 100 mL of medium was added in 150 mL conical flasks, inoculated with a ring of target strain, and then incubated for 48 h at 30°C (yeast) and 24 h at 37°C (LAB). The microscopy was used to continuously count until obtained 10^8^ CFU/mL seed fermentation broth. Solid-state fermentation. Sorghum (400 g) was added to 500 mL of water in the 3000 L beaker, and mixed the liquefied enzyme (10 U/g) in boiling water (100°C) for 2 h, and then added glucoamylase (50 u/g) maintaining 4 h at 60°C. Reducing sugar of the sorghum extracts about 50 ~ 90 g/kg. The beaker autoclaved at 115°C for 15 min. After cooling, seed fermentation broth was added in the beaker with the cellular population of 1×10^5^ CFU/g wet sorghum, and then experiments were carried out in 150 mL conical flasks which contained 100 g of sorghum. The flasks were then sealed and incubated at 30°C. In order not to interrupt the fermentation process, 30 flasks were used to fermented according to the above experimental conditions, and three flasks were randomly selected from the same fermentation conditions at 1, 2, 3, 4, 5, 10, 15, 20, 25 and 28 days respectively. After fermentation, the sorghums were used enumeration of different strains, and the rest withdrew and stored at - 20°C for analysis of environmental factors and flavor compounds.

### Enumeration of different strains

After fermentation, 10 g sorghums were added to 25 mL phosphate buffer saline (PBS, 0.01M, pH7.2), vortex mix 3000 rpm for 30 s (Dragonlab MX-E, Beijing, China), and under 4 °C for 30 min. The supernatant was gradient diluted and spread plate. Four kinds of yeasts enumeration were carried out on Wallerstein Laboratory nutrient (WLN) medium (78), in which the strains showed different macroscopic features (texture, surface, margin, and color),. *Lactobacillus* enumeration was carried out on MRS Broth (DE MAN, ROGOSA, SHARPE) (34). Standard deviations were calculated from triplicate repetitions of the enumeration.

### Statistical analysis

Standard statistical analyses were conducted with XLSTAT (v.19.02.42992). Heatmap, Variation partitioning analysis, Redundancy analysis (RDA), the Monte Carlo permutation test was calculated by the R program (v. 3.4.0). In the Heatmap, flavor compounds were transformed by z-score. Clustering analysis was performed using the Pearson correlation coefficient, and Euclidean distance based on the flavor compounds content during the fermentation process. The Variation partitioning analysis resulted in five environmental factors and five microbes’ average abundance. In constrained ordination, representational difference analysis (RDA) was used to identify the relationship of samples, environmental factors and microbes. The Monte Carlo permutation test was used to examine the significance of the correlation between environmental factors and species distribution. All the analyses were performed using functions in the Vegan package (v. 2.4-3) (79). The Spearman correlation coefficient (*ρ*) and Paired-sample t-test were calculated with SPSS Statistics 22, in which *ρ* > 0.6 and *ρ* > 0.8 were representing strongly and highly correlated. The visualization objects of interaction of flavor compounds and microbes and co-occurring analysis were drawn with Gephi (v. 0.9.1) (22).

## SUPPLEMENTAL MATERIAL

SUPPLEMENTAL FILE S1.

DATASET S1, XLSX file.

DATASET S2, XLSX file.

DATASET S3, XLSX file.

## ACKNOWLEDGMENTS

We would like to thank Shanxi Xinghuacun Fenjiu Distillery Co., Ltd. for the samples, Peng Wang and Jianchun Lin for statistical methods and sample collection.

## FUNDING

This work was supported by the National Natural Science Foundation of China (NSFC) (31530055), National Key R&D Program of China (2018YFD0400402, 2016YFD0400500), Jiangsu Province Science and Technology Project (BE2017705), China Postdoctoral Science Foundation (2017M611702), the Postgraduate Research & Practice Innovation Program of Jiangsu Provence (KYCX18_1798), and national first-class discipline program of Light Industry Technology and Engineering (LITE2018-12), the Priority Academic Program Development of Jiangsu Higher Education Institutions, the 111 Project (111-2-06), the Collaborative Innovation Center of Jiangsu Modern Industrial Fermentation.

The authors declare that there are no conflicts of interest.

## REFERENCES

1. Smid EJ, Lacroix C. 2013. Microbe-microbe interactions in mixed culture food fermentations. Curr Opin Biotech 24:148–154.

2. Bokulich NA, Thorngate JH, Richardson PM, Mills DA. 2014. Microbial biogeography of wine grapes is conditioned by cultivar, vintage, and climate. PNAS 111:139–148.

3. Meersman E, Steensels J, Mathawan M, Wittocx PJ, Saels V, Struyf N, Bernaert H, Vrancken G, Verstrepen KJ. 2013. Detailed analysis of the microbial population in malaysian spontaneous cocoa pulp fermentations reveals a core and variable microbiota. PLoS ONE 8:e81559.

4. Rodríguez-Lerma GK, Gutiérrez-Moreno K, Cárdenas-Manríquez M, Botello-Álvarez E, Jiménez-Islas H, Rico-Martínez R, Navarrete-Bolaños JL. 2011. Microbial ecology studies of spontaneous fermentation: starter culture selection for prickly pear wine production. J Food Sci Technol 76:M346–M352.

5. Holzapfel W. 2002. Appropriate starter culture technologies for small-scale fermentation in developing countries. Int J Food Microbiol 75:197–212.

6. Giraffa G. 2004. Studying the dynamics of microbial populations during food fermentation. FEMS Microbiolo Rev 28:251–260.

7. Wang ZM, Lu ZM, Shi JS, Xu ZH. 2016. Exploring flavour-producing core microbiota in multispecies solid-state fermentation of traditional Chinese vinegar. Sci Rep 6:26818.

8. Sieuwerts S, de Bok FA, Hugenholtz J, Je VHV. 2008. Unraveling microbial interactions in food fermentations: from classical to genomics approaches. Appl Environ Microbiol 74:4997–5007.

9. Wolfe BE, Button JE, Santarelli M, Dutton RJ. 2014. Cheese rind communities provide tractable systems for in situ and in vitro studies of microbial diversity. Cell 158:422–433.

10. De Filippis F, La Storia A, Stellato G, Gatti M, Ercolini D. 2014. A selected core microbiome drives the early stages of three popular Italian cheese manufactures. PLoS ONE 9:e89680.

11. Wolfe BE, Dutton RJ. 2015. Fermented foods as experimentally tractable microbial ecosystems. Cell 161:49–55.

12. Hu XL, Du H, Ren C, Yan X. 2016. Illuminating anaerobic microbial community and cooccurrence patterns across a quality gradient in Chinese liquor fermentation pit muds. Appl Environ Microbiol 82:2506–2515.

13. Wu Q, Ling J, Xu Y. 2014. Starter culture selection for making Chinese sesame-flavored liquor based on microbial metabolic activity in mixed-culture fermentation. Appl Environ Microbiol 80:4450–4459.

14. Armstrong MS, Boyd CE, Lovell RT. 1986. Environmental factors affecting flavor of channel catfish from production ponds. The Progressive Fish-Culturist 48:113–119.

15. Pothakos V, Illeghems K, Laureys D, Spitaels F, Vandamme P, De Vuyst L. 2016. Acetic acid bacteria in fermented food and beverage ecosystems, p 73–99, Acetic Acid Bacteria. Springer.

16. Liu J, Wu Q, Wang P, Lin JC, Huang L, Xu Y. 2017. Synergistic effect in core microbiota associated with sulfur metabolism in spontaneous Chinese liquor fermentation. Appl Environ Microbiol 83:e01475–17.

17. Cardinale M, Grube M, Erlacher A, Quehenberger J, Berg G. 2015. Bacterial networks and co-occurrence relationships in the lettuce root microbiota. Environ Microbiol 17:239–252.

18. Parente E, Cocolin L, De Filippis F, Zotta T, Ferrocino I, O’Sullivan O, Neviani E, De Angelis M, Cotter PD, Ercolini D. 2016. FoodMicrobionet: A database for the visualisation and exploration of food bacterial communities based on network analysis. Int J Food Microbiol 219:28–37.

19. Jin GY, Zhu Y, Xu Y. 2017. Mystery behind Chinese liquor fermentation. Trends Food Sci Tech 63:18–28.

20. Zhang CL, Ao ZH, Chui WQ, Shen CH, Tao WY, Zhang SY. 2012. Characterization of the aroma-active compounds in *Daqu*: a tradition Chinese liquor starter. Eur Food Res Technol 234:69–76.

21. Lemos LN, Fulthorpe RR, Triplett EW, Roesch LF. 2011. Rethinking microbial diversity analysis in the high throughput sequencing era. J Microbiol Methods 86:42–51.

22. Wang X, Du H, Xu Y. 2017. Source tracking of prokaryotic communities in fermented grain of Chinese strong-flavor liquor. Int J Food Microbiol 244:27–35.

23. Rui JP, Li JB, Zhang SH, Yan XF, Wang YP, Li XZ. 2015. The core populations and co-occurrence patterns of prokaryotic communities in household biogas digesters. Biotechnol Biofuels 8:158–173.

24. Gao WJ, Fan WL, Xu Y. 2014. Characterization of the key odorants in light aroma type Chinese liquor by gas chromatography–olfactometry, quantitative measurements, aroma recombination, and omission studies. J Agric Food Chem 62:5796–5804.

25. Cao Y, Hu Y. 2010. Variation of Aromatic Components in Solid Phase Fermented Grains During Fermentation of Fen Liquor. Food Sci 31:367–371.

26. Barberan A, Bates ST, Casamayor EO, Fierer N. 2012. Using network analysis to explore co-occurrence patterns in soil microbial communities. ISME J 6:343–351.

27. De Filippis F, Genovese A, Ferranti P, Gilbert JA, Ercolini D. 2016. Metatranscriptomics reveals temperature-driven functional changes in microbiome impacting cheese maturation rate. Sci Rep 6:21871.

28. Steinhauser D, Krall L, Müssig C, Büssis D, Usadel B. 2007. Correlation Networks, Analysis of Biological Networks. John Wiley & Sons, Inc.

29. Yang J, Leskovec J. 2014. Overlapping communities explain core–periphery organization of networks. P IEEE 102:1892–1902.

30. Ercolini D, Pontonio E, Filippis FD, Minervini F, Storia AL, Gobbetti M, Cagno RD. 2013. Microbial ecology dynamics during rye and wheat sourdough preparation. Appl Environ Microbiol 79:7827–7836.

31. Yang Y, Gao Y, Wang S, Xu D, Yu H, Wu L, Lin Q, Hu Y, Li X, He Z. 2014. The microbial gene diversity along an elevation gradient of the Tibetan grassland. ISME J 8:430.

32. De Bruyn F, Zhang SJ, Pothakos V, Torres J, Lambot C, Moroni AV, Callanan M, Sybesma W, Weckx S, De Vuyst L. 2017. Exploring the impacts of postharvest processing on the microbiota and metabolite profiles during green coffee bean production. Appl Environ Microbiol 83:e02398–16.

33. Tang J, Tang X, Tang M, Zhang X, Xu X, Yi Y. 2017. Analysis of the Bacterial Communities in Two Liquors of Soy Sauce Aroma as Revealed by High-Throughput Sequencing of the 16S rRNA V4 Hypervariable Region. Biomed Res Int 2017:1–9.

34. Yang X, Teng K, Jie Z, Wang F, Tong Z, Ai G, Han P, Bai F, Jin Z. 2017. Transcriptome responses of Lactobacillus acetotolerans F28 to a short and long term ethanol stress. Sci Rep 7:2650.

35. Toh H, Morita H, Tsuji H, Iwashita K, Goto N, Nakayama J, Sekine M, Kato Y, Suzuki K, Fujita N. 2015. Complete genome sequence of Lactobacillus acetotolerans RIB 9124 (NBRC 13120) isolated from putrefied (hiochi) Japanese sake. J Biotechnol 214:214–215.

36. Wu Q, Zhu W, Wang W, Xu Y. 2015. Effect of yeast species on the terpenoids profile of Chinese light-style liquor. Food Chem 168:390–395.

37. Tian X, Cui H, Wang X. 2016. Research Progress on the Application of the Fermentation Products of Geotrichum candidum. Anhui Agr Sci Bull 22:31–32.

38. Noronhadacosta P, Spencermartins I, Loureiro V, Rodrigues C. 2010. Fatty acid patterns of film‐forming yeasts and new evidence for the heterogeneity of Pichia membranaefaciens. Lett Appl Microbiol 23:79–84.

39. Gong GL, Ma LY, Chen XF. 2014. Isolation and improvement of Saccharomyces cerevisiae for producing the distilled liquor. J Chem Pharm Res 6:283–288.

40. Kong Y, Wu Q, Zhang Y, Xu Y. 2014. *In situ* analysis of metabolic characteristics reveals the key yeast in the spontaneous and solid-state fermentation process of Chinese light-style liquor. Appl Environ Microbiol 80:3667–3676.

41. Hu XL, Wang HY, Wu Q, Xu Y. 2014. Development, validation and application of specific primers for analyzing the clostridial diversity in dark fermentation pit mud by PCR-DGGE. Bioresource Technol 163:40–47.

42. Wang P, Wu Q, Jiang X, Wang Z, Tang J, Xu Y. 2017. *Bacillus licheniformis* affects the microbial community and metabolic profile in the spontaneous fermentation of *Daqu* starter for Chinese liquor making. Int J Food Microbiol 250:59–67.

43. Gao W, Fan W, Xu Y. 2014. Characterization of the Key Odorants in Light Aroma Type Chinese Liquor by Gas Chromatography–Olfactometry, Quantitative Measurements, Aroma Recombination, and Omission Studies. Agric Food Chem 62:5796–5804.

44. Niu Y, Yao Z, Xiao Q, Xiao Z, Ma N, Zhu J. 2017. Characterization of the key aroma compounds in different light aroma type Chinese liquors by GC-olfactometry, GC-FPD, quantitative measurements, and aroma recombination. Food Chem 233:204–215.

45. Monnet C, Dugat-Bony E, Swennen D, Beckerich J-M, Irlinger F, Fraud S, Bonnarme P. 2016. Investigation of the activity of the microorganisms in a reblochon-style Cheese by metatranscriptomic analysis. Front Microbiol 7:536.

46. Li S, Li P, Liu X, Luo LX, Lin WF. 2016. Bacterial dynamics and metabolite changes in solid-state acetic acid fermentation of Shanxi aged vinegar. Appl Environ Microbiol 100:4395–4411.

47. Kable ME, Srisengfa Y, Laird M, Zaragoza J, McLeod J, Heidenreich J, Marco ML. 2016. The core and seasonal microbiota of raw bovine milk in tanker trucks and the impact of transfer to a milk processing facility. Mbio 7:e00836–16.

48. Annan NT, Poll L, Sefa-Dedeh S, Plahar WA, Jakobsen M. 2003. Volatile compounds produced by *Lactobacillus fermentum*, *Saccharomyces cerevisiae* and *Candida krusei* in single starter culture fermentations of Ghanaian maize dough. J Appl Microbiol 94:462–474.

49. Laakso K, Koskenniemi K, Koponen J, Kankainen M, Surakka A, Salusjärvi T, Auvinen P, Savijoki K, Nyman TA, Kalkkinen N. 2011. Growth phase-associated changes in the proteome and transcriptome of *Lactobacillus rhamnosus* GG in industrial-type whey medium. Microb Biotechnol 4:746–766.

50. Wang PX, Mao J, Meng XY, Li XZ, Liu YY, Feng H. 2014. Changes in flavour characteristics and bacterial diversity during the traditional fermentation of Chinese rice wines from Shaoxing region. Food Control 44:58–63.

51. Li P, Li S, Cheng L, Luo L. 2014. Analyzing the relation between the microbial diversity of *Daqu* and the turbidity spoilage of traditional Chinese vinegar. Appl Environ Microbiol 98:6073–84.

52. Almeida M, Hébert A, Abraham A-L, Rasmussen S, Monnet C, Pons N, Delbès C, Loux V, Batto J-M, Leonard P, Kennedy S, Ehrlich SD, Pop M, Montel M-C, Irlinger F, Renault P. 2014. Construction of a dairy microbial genome catalog opens new perspectives for the metagenomic analysis of dairy fermented products. BMC Genomics 15:1101–1116.

53. Balat M. 2011. Production of bioethanol from lignocellulosic materials via the biochemical pathway: A review. Energ Convers Manage 52:858–875.

54. Mattanovich D, Sauer M, Gasser B. 2016. Industrial microorganisms: Pichia pastoris, p 687–714, Ind Biotechnol, Wiley-Blackwell.

55. Saerens S, Swiegers JH. 2017. Production of low-alcohol or alcohol-free beer with *Pichia kluyveri* yeast strains. Google Patents.

56. Rojas V, Gil JV, Piñaga F, Manzanares P. 2001. Studies on acetate ester production by non-*Saccharomyces* wine yeasts. Int J Food Microbiol 70:283–289.

57. Veeraragavan K, Colpitts T, Gibbs B. 1990. Purification and Characterization of 2 Distinct Lipases from Geotrichum-Candidum. J Am Ceram Soc 95:2140–2147.

58. Burkert JF, Maugeri F, Rodrigues MI. 2004. Optimization of extracellular lipase production by *Geotrichum* sp. using factorial design. Bioresource Technol 91:77–84.

59. Sukpipat W, Komeda H, Prasertsan P, Asano Y. 2017. Purification and characterization of xylitol dehydrogenase with l-arabitol dehydrogenase activity from the newly isolated pentose-fermenting yeast *Meyerozyma caribbica* 5XY2. J Biosci Bioeng 123:20–27.

60. Bourrie BC, Willing BP, Cotter PD. 2016. The microbiota and health promoting characteristics of the fermented beverage kefir. Front Microbiol 7:674.

61. Gethins L, Guneser O, Demirkol A, Rea MC, Stanton C, Ross RP, Yuceer Y, Morrissey JP. 2015. Influence of carbon and nitrogen source on production of volatile fragrance and flavour metabolites by the yeast Kluyveromyces marxianus. Yeast 32:67–76.

62. Ren K, Lamsal BP. 2017. Synthesis of some glucose-fatty acid esters by lipase from Candida antarctica and their emulsion functions. Food Chem 214:556–563.

63. Ivić JT, Milosavić N, Dimitrijević A, Jankulović MG, Bezbradica D, Kolarski D, Veličković D. 2017. Synthesis of medium-chain length capsinoids from coconut oil catalyzed by *Candida rugosa* lipases. Food Chem 218:505–508.

64. He Y, Li J, Kodali S, Chen B, Guo Z. 2017. Rationale behind the near-ideal catalysis of Candida antarctica lipase A (CAL-A) for highly concentrating ω-3 polyunsaturated fatty acids into monoacylglycerols. Food Chem 219:230–239.

65. Englezos V, Torchio F, Cravero F, Marengo F, Giacosa S, Gerbi V, Rantsiou K, Rolle L, Cocolin L. 2016. Aroma profile and composition of Barbera wines obtained by mixed fermentations of *Starmerella bacillaris* (synonym *Candida zemplinina*) and *Saccharomyces cerevisiae*. LWT-Food Sci Technol 73:567–575.

66. Zhang XL, Tian XQ, Ma LY, Feng B, Liu QH, Yuan LD, Fan CQ, Huang HL, Yang Q. 2015. Biodiversity of the symbiotic bacteria associated with toxic marine dinoflagellate *Alexandrium tamarense*. JBM 3:23–28.

67. Toju H, Tanabe AS, Yamamoto S, Sato H. 2012. High-coverage ITS primers for the DNA-based identification of ascomycetes and basidiomycetes in environmental samples. PLoS ONE 7:e40863.

68. Ren G, Ren W, Teng Y, Li Z. 2015. Evident bacterial community changes but only slight degradation when polluted with pyrene in a red soil. Front Microbiol 6:22.

69. Gabert J, Beillard E, Vh VDV, Bi W, Grimwade D, Pallisgaard N, Barbany G, Cazzaniga G, Cayuela JM, Cavé H. 2006. Real-time quantitative PCR (qPCR) and reverse transcription-qPCR for detection and enumeration of total yeasts in wine. Appl Environ Microbiol 72:7148–7155.

70. Mayrhofer S, Filipp R, Lehner D, Reiterich C, Kneifel W, Domig KJ. 2016. Suitability of different PCR-DGGE primer sets for the monitoring of lactic acid bacteria in wine. S Afr J Enol Vitic 35:185–195.

71. Caporaso JG, Kuczynski J, Stombaugh J, Bittinger K, Bushman FD, Costello EK, Fierer N, Pena AG, Goodrich JK, Gordon JI, Huttley GA, Kelley ST, Knights D, Koenig JE, Ley RE, Lozupone CA, McDonald D, Muegge BD, Pirrung M, Reeder J, Sevinsky JR, Turnbaugh PJ, Walters WA, Widmann J, Yatsunenko T, Zaneveld J, Knight R. 2010. QIIME allows analysis of high-throughput community sequencing data. Nat Methods 7:335–336.

72. Edgar RC, Haas BJ, Clemente JC, Quince C, Knight R. 2011. UCHIME improves sensitivity and speed of chimera detection. Bioinformatics 27:2194–2200.

73. Edgar RC. 2010. Search and clustering orders of magnitude faster than BLAST. Bioinformatics 26:2460–2461.

74. Tao Y, Li J, Rui J, Xu Z, Zhou Y, Hu X, Wang X, Liu M, Li D, Li X. 2014. Prokaryotic communities in pit mud from different-aged cellars used for the production of Chinese strong-flavored liquor. Appl Environ Microbiol 80:2254–2260.

75. Edgar RC. 2013. UPARSE: Highly accurate OTU sequences from microbial amplicon reads. Nat Methods 10:996–998.

76. Li P, Lin W, Liu X, Wang X, Gan X, Luo L, Lin WT. 2017. Effect of bioaugmented inoculation on microbiota dynamics during solid-state fermentation of *Daqu* starter using autochthonous of *Bacillus*, *Pediococcus*, *Wickerhamomyces* and *Saccharomycopsis*. Food Microbiol 61:83–92.

77. Wu Q, Chen L, Xu Y. 2013. Yeast community associated with the solid state fermentation of traditional Chinese *Maotai*-flavor liquor. Int J Food Microbiol 166:323–330.

78. Wu Q, Kong Y, Xu Y. 2015. Flavor profile of Chinese liquor is altered by interactions of intrinsic and extrinsic microbes. Appl Environ Microbiol 82:422–430.

79. Oksanen J, Kindt R, Legendre P, O’Hara B, Stevens MHH, Oksanen MJ, Suggests M. 2008. The vegan package, http://vegan.r-forge.r-project.org/.

